# CMOT: Cross Modality Optimal Transport for multimodal inference

**DOI:** 10.1101/2022.10.15.512387

**Authors:** Sayali Alatkar, Daifeng Wang

## Abstract

Biological mechanisms are complex spanning multiple facets, each providing a unique view of the underlying mechanism. Recent single-cell technologies such as scRNA-seq and scATAC-seq have facilitated parallel probing into each of these facets, providing multimodal measurements of individual cells. An integrative analysis of these single-cell modalities can therefore provide a comprehensive understanding of specific cellular and molecular mechanisms. Despite the technological advances, simultaneous profiling of multiple modalities of single cells continues to be challenging as opposed to single modality measurements, and therefore, it may not always be feasible to conduct such profiling experiments (e.g., missing modalities). Furthermore, even with the available multimodalities, data integration remains elusive since modalities may not always have paired samples, leaving partial to no correspondence information.

To address those challenges, we developed Cross-Modality Optimal Transport (CMOT), a computational approach to infer missing modalities of single cells based on optimal transport (OT). CMOT first aligns a group of cells (source) within available multi-modal data onto a common latent space. Then, it applies optimal transport to map the cells with a single modality (target) to the aligned cells in source from the same modality by minimizing their cost of transportation using Wasserstein distance. This distance can be regularized by prior knowledge (e.g., cell types) or induced cells clusters to improve mapping, and an entropy of transport to speed up OT computations. Once transported, CMOT uses K-Nearest-Neighbors to infer the missing modality for the cells in target from another modality of mapped cells in source. Furthermore, the alignment of CMOT works for partially corresponding information from multi-modalities, i.e., allowing a fraction of unmatched cells in source. We evaluated CMOT on several emerging multi-modal datasets, e.g., gene and protein expression and chromatin accessibility in developing brain, cancers, and immunology. We found that not only does CMOT outperform existing state-of-art methods, but its inferred gene expression is biologically interpretable such as classifying cell types and cancer types. Finally, CMOT is open source and available on our github: https://github.com/daifengwanglab/CMOT

## 1 Introduction

Single-cell sequencing technologies can measure different characteristics of single cells across multi-omics such as the genomics, transcriptomics, epigenomics, proteomics. Such high-resolution measurements have enabled exploring individual cells to reveal cellular and molecular mechanisms and study cell-to-cell functional variations. For example, scCAT-seq, sci-CAR, and 10xMultiome measure single-cell gene expression and chromatin accessibility [1], [2], and CITE-seq measures gene and protein expression of single cells [3], [4]. However, simultaneous profiling of such multi-omics and additional modalities continues to be a challenging task especially because of high sequencing costs, low recovery of individual cells, and sparse and noisy data [5]. Owing to these challenges, single-cell multimodal data generation may not always be feasible. This leads to the question of how we can use available multi-modalities to infer missing modalities.

Several prior works have tackled modality inference. Seurat [6] infers the missing modality of a cell by weighting nearest neighboring cells with multimodalities available. MOFA+ [7] uses Bayesian factor analysis to identify a lower dimensional representation of the data to infer the missing modality. However, they only work with multimodal data that must come from the same cells (i.e., fully corresponding). Alignment-based methods like non-linear manifold alignment [8] have shown to align multimodalities with partial cell-to-cell correspondence information but not been extended to cross-modality inference.

Machine learning has also emerged to help modality inference. For instance, TotalVI [4] builds a variational autoencoder that infers missing protein profiles from gene expression using CITE-seq data. Babel [9] uses two autoencoders, one for each modality to infer a joint latent representation for cross-modal translation. Polarbear [10] also uses autoencoders, however, trains on both single and multi-modal data to infer each modality. However, such autoencoder-based approaches are unsupervised, and therefore lack a mechanism to introduce prior knowledge about underlying data distribution. Moreover, training autoencoders typically requires considerable amounts of data and time with intensive hyperparameter tuning.

Optimal Transport (OT), an efficient approach uses prior knowledge about data distribution to find an optimal mapping between the distributions [11]. OT can also work on small datasets with limited parameters. Recently, OT has been applied to single-cell multiomics data for various applications [12]–[15]. Schiebinger et. al. [12] used OT to model the developmental trajectory of single-cell gene expression. through unbalanced optimal transport. Single-cell integrative analysis frameworks like SCOT [13], SCOTv2 [14], and Pamona [15] further extended the original OT problem for multi-omics data alignment. Another work [16] used OT with an additional entropic regularization term to improve the unsupervised clustering of single-cell data to understand cell types and cellular states better. However, OT has not yet been applied for cross-modality inference. Thus, we propose that integrating OT with multimodal data alignment can work for cross-modality inference and address the above limitations of prior works.

Particularly, we developed CMOT (Cross-Modality Optimal Transport), a computational approach to infer missing modalities of single cells. CMOT first aligns the cells with multimodal data (source), and then applies OT to map the cells from single modality (target) to the source cells via shared modality. Finally, CMOT uses the K-Nearest-Neighbors (KNN) of source cells to infer missing modality for target cells. Moreover, CMOT does not need paired multi-modal data for alignment. We found that not only does CMOT outperform existing state-of-art methods, but its inferred gene expression is biologically interpretable by evaluating on emerging single-cell multi-omics datasets. Finally, CMOT is open source at: https://github.com/daifengwanglab/CMOT.

## 2 Methods and Materials

### 2.1 Overview of Cross-Modality Optimal Transport

CMOT (Cross-Modality Optimal Transport) is a computational approach for cross-modality inference of single cells. As shown in **Fig. 1**, CMOT has three main steps: **Step A** –Manifold alignment to project the cells with available multimodal data (source cells) onto a common low-dimensional latent space. The cells from different modalities can be unpaired or have partial correspondence; **Step B** – Optimal Transport to map the cells with a single modality (target cells) to the aligned source cells from the same modality. To this end, we minimize the Wasserstein distance between the source and target cells and can further regularize the minimization using prior knowledge (e.g., cell types) or induced cell clusters and entropy of transport; **Step C** – K-Nearest-Neighbors to infer the missing or unprofiled modality of target cells using another modality of nearest mapped source cells.

**Figure 1:**
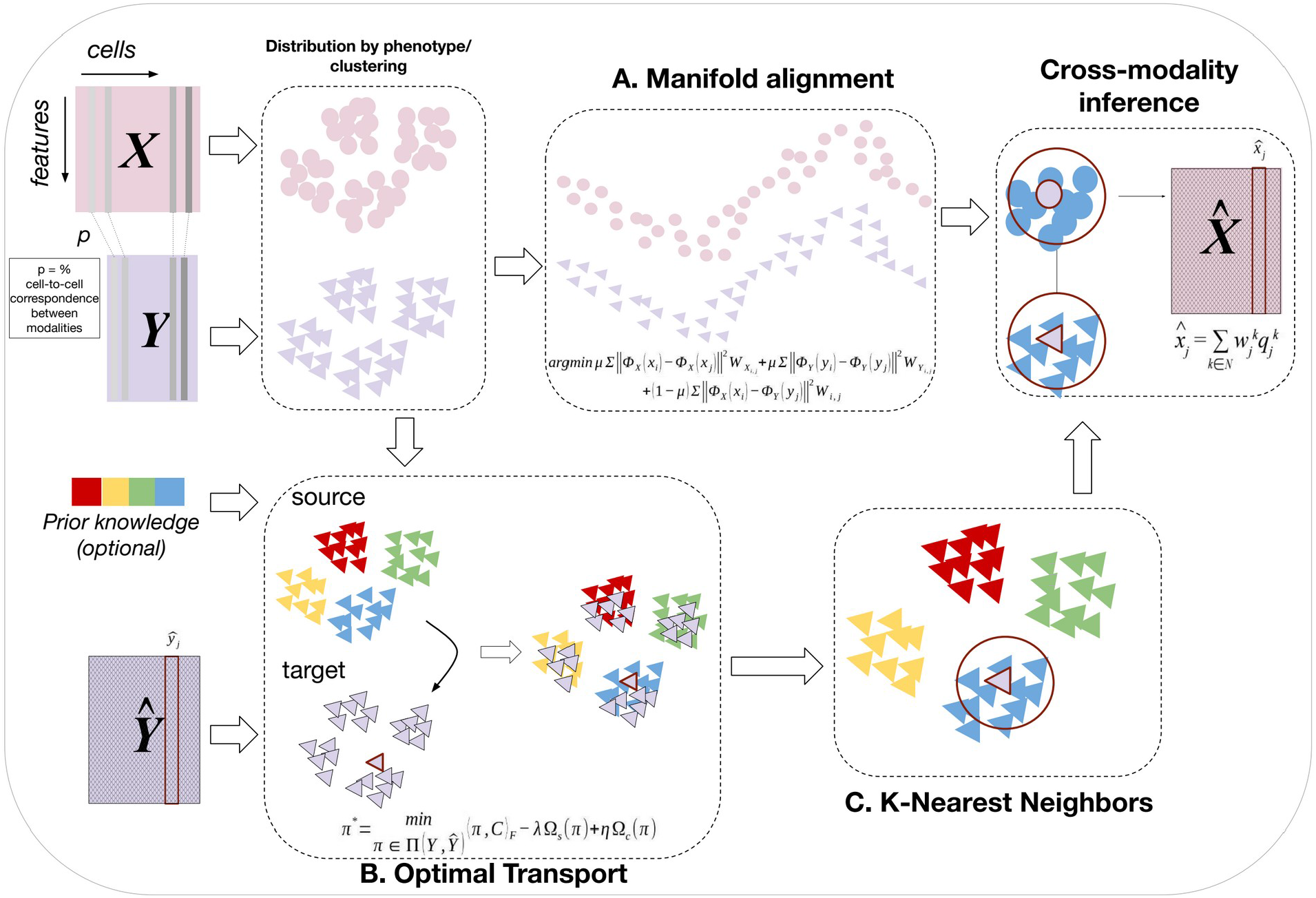
Cross-Modality Optimal Transport (CMOT): CMOT has three main steps: (**A**) Manifold Alignment, (**B**) Optimal Transport, and (**C**) K-Nearest Neighbors inference. CMOT inputs two multimodalities *X* and *Y* (source), where the cells in *X* and *Y* need not be completely corresponding. The cell-to-cell correspondence information between *X* and *Y* can be specified through *p.* CMOT aligns *X* and *Y* using non-linear manifold alignment (NMA) onto a common low-dimensional latent space. Then, CMOT uses optimal transport (OT) to map the aligned cells in source *Y* to the cells in target 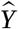, where *Y* and *Y* share modalities. CMOT minimizes the cost of transportation by finding the Wasserstein distance between cells in *Y* and *Y* which is further regularized by prior knowledge or induced cell clusters and entropy of transport. Finally, CMOT infers the missing modality 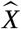 for cells in *Y* using K-nearest neighbors (KNN). It calculates a weighted average of the k-nearest mapped cells in *Y* for every cell in *Y*, using their values from *X*, and infers 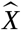.

#### Step A: Manifold alignment to align source cells with multimodal data

In this step, we align the multimodalities using Nonlinear Manifold Alignment (NMA). Alignment is an important step that accounts for when the source cells have partial correspondence. NMA is based on a manifold hypothesis that high dimensional multimodal datasets have similar underlying low dimensional manifolds, and therefore, they can be projected onto a common manifold space that preserves the local geometry of each modality and minimizes the differences between the manifolds of modalities. We define *X* = {*x_i_*}_*i*,…,*s_X_*_ and *Y* = {*y_j_*}_*j*= 1,…,*s_Y_*_ as two multimodal measurements of *s_X_* source cells in Modality *X* and *s_Y_* source cells in Modality *Y*, where *x_i_* ∈ *R^d_X_^* and *y_k_* ∈ *R^d_Y_^* represent the measurements of *d_X_* features in *i*^th^ cell in Modality *X*, and *d_Y_* features in *j*^th^ cell in Modality *Y*, respectively. We also define *W_x_* ∈ *R*^*s_X_* × *s_X_*^ and *W_y_* ∈ *R*^*s_Y_* × *s_Y_*^ as cell similarity matrices for *X* and *Y*, respectively, where each similarity matrix is constructed by connecting a cell with its *K* nearest neighboring cells within the modality. The partial prior known cell-to-cell correspondence information can be quantified by *p* (0 < *p*<100) using cross-modal similarity matrix *W* ∈ *R*^*s_x_* × *s_X_*^, i.e., *p*% of paired cells across modalities. NMA then learns two mapping functions *Φ_X_* and *Φ_Y_* that project *x_i_* and *y_j_* to *Φ_X_* (*x_i_*) ∈ *R^d^* and *Φ_Y_*(*y_j_*) ∈ *R^d^,* respectively onto a common manifold space with dimension *d* ≪ *min*(*d_x_, d_y_*). The *d*-dimensional manifold preserves the local geometry of each modality and minimizes the distances between corresponding samples after projection. Solving manifold alignment can be reformulated as manifold co-regularization in reproducing kernel Hilbert spaces. The manifold alignment optimization term is:

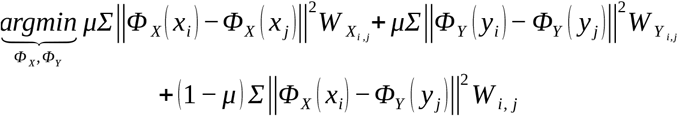
where the first two terms preserve the local geometry within each modality, the similarity matrices *W_x_* ∈ *R*^*s_X_* × *s_X_*^ and *W_y_* ∈ *R*^*s_Y_* × *s_Y_*^ model the relationships of the cells in each modality that can be identified by *k*-nearest neighbor graph, and the third term preserves the correspondence information across *X* and *Y* modeled by *W* ∈ *R*^*s_X_* × *s_Y_*^. The parameter *μ* controls the trade-off between conserving the local geometry of each modality and cell-to-cell correspondences across modalities.

Also, two modalities are not required to have a complete correspondence between the cells. Therefore, *W* is a binary correspondence matrix between cells of *X* and *Y* such that if *p*=1, i.e., 100% correspondence across cells in *X* and *Y*, *W* would be an identity matrix. For *p*<1, *W_i,j_*=1 if *x_i_* and *y_j_* are the corresponding cells and 0 otherwise. After alignment, the resulting *d*-dimensional modalities share a common latent space that can easily be compared using Euclidean distances. For instance, for every cell *y_j_* ∈ *Y*, we find an aligned cell *X_j,a_* ∈ *X* by finding the closest cell in *X* using the Euclidean distance. To implement this alignment step, we used the Python functions in ManiNetCluster [17].

#### Step B: Optimal Transport to map source and target cells by shared modality

The optimal transport theory [18], [19] tries to find the most efficient mapping *π*\* that transports one probability distribution to another with minimum transportation cost. A mapping 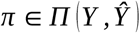 represents transport plan to map cells from source (*Y*) and target 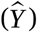 modalites, where 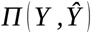 contains all probabilistic mappings between the source (*Y*) and target cells 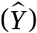. The classical OT distance (Wasserstein distance) gives the distance between two probability distributions as the transportation cost. *C* is cost matrix where 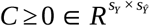 and *C*(*i,j*) represents the pairwise cost of mapping the source cell *y_i_* ∈ *Y*to the target cell 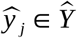. For discrete probability distributions like *Y* and 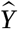 over the same metric spaces (i.e., matched features of shared modality), we define the OT problem as:

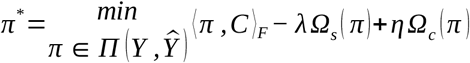
where the first term computes the Frobenius dot product 〈.,.〉_*F*_ between the cost matrix *C* and *π*. The set *Π* is defined as 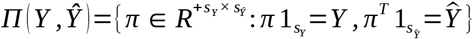.

The second term, also called entropic regularization, calculates the entropy of transportation for *π* where 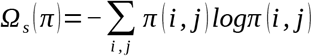. Entropic regularization addresses the computational complexity of OT as the sample size increases [15]. The intuition behind this term is to relax the sparsity constraints of the OT problem by increasing its entropy so that *π*\* is denser, as source cells (*Y*) are distributed more towards target cells 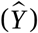. The resulting formulation is strictly convex and can be solved through Sinkhorn’s Algorithm [20]. As the parameter increases, the sparsity of *π*\* decreases, giving a smoother transport. The parameter *λ* weights the entropic regularization.

The third term is the label regularizer [21], 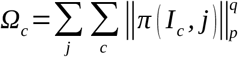, where *I_c_* contains the index of rows in *π* related to the source cells (*Y*) that belong to class *c* if we have such prior knowledge, e.g., known cell types. Hence, *π* (*I_C_, j*) is a vector containing the coefficients of the *j*th column of *π* associated with class *c* where the *j*th column in *π*(*I_C_, j*) represents the *j*th target cell in 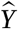. The norm 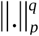. denotes the *l_p_* norm to the power of *q*. The parameter *η* weights the label regularization. The intuition behind this term is to penalize the mappings that match together samples from different labels. This means that even if we do not have the label information for the target cells 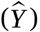, we can promote group sparsity within the columns of *π* such that each target cell is only associated with a class. However, in the absence of such label information, we can compute our own labels through unsupervised clustering techniques like hierarchical clustering to induce cell clusters as labels for source cells in *Y*. Finally, to map the source cells (*Y*) to the target space 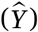, we use barycentric mapping using *Y*^(*t*)^ = *s_x_π* * *Y* [21]. Now, we can easily compare *Y*^(*t*)^ and 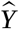 using euclidean distance. To solve the OT optimization, Python optimal transport (POT) package is used [22].

#### Step C: K-Nearest Neighbors to infer the additional modality of target cells

Finally, we apply K-Nearest Neighbors (KNNs) to infer the missing modality 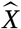 of target cells. For each target cell 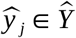, we find its KNN in *Y*^(*t*)^ using Euclidean distance. Let *S_j_*={*c_j,k_*: *k* =1,2,..*K*} be the set of k nearest neighboring cells of 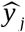 in *Y*^(*t*)^, where *c_j,k_* is the *k*^th^ cell in *Y*^(*t*)^. For cells in *S_j_*, we use their values from the aligned modality 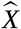 to define set 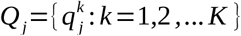, where 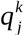 represents the profile of the cell *c_j,k_* within the aligned modality *X*. Finally we calculate the weighted average of all cells in *Q_j_* to get 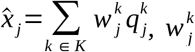 is the weightage given to 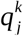 such that 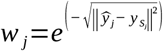. Thus, we get the corresponding modality 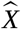 for 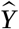. We used sklearn’s [23] nearest neighbor function for KNN implementation.

### 2.2 Single-cell multi-omics datasets

We tested CMOT on four single-cell multiomics datasets: (1) Gene expression and chromatin accessibility of single cells in human and mouse brains (scRNA-seq & scATAC-seq) [1], [24]; (2) Gene and protein expression of peripheral blood mononuclear cells (CITE-seq) [4]; (3) Gene expression and chromatin accessibility of A549 lung cancer cells (sci-CAR) [2]; (4) Gene expression and chromatin accessibility of pancancer cells (scCAT-seq) [25]. All details on data and data processing are available in supplementary methods.

### 2.3 Parameter selection

In Step A, we found the optimal alignment by testing different values of *d* common manifold dimensions and *k* nearest neighbors for building the similarity within each modality (**Fig. S5**). In Step B and Step C, we performed cross-validation to select the regularization coefficients *λ* and *η* for optimal transport and *k*-nearest neighbors for modality inference. Also, for all datasets applied in this paper, we held out a 20% testing set to report CMOT’s performance. For datasets with no prior knowledge (e.g., cell types), we induced cell labels by cell clusters through hierarchical clustering of the training set, when training the final model (see Supplementary Methods). We split the training data into training and validation sets to select parameters through 5-fold cross-validation (see Supplementary Methods).

### 2.4 Evaluation

#### Inference versus measurement

To evaluate CMOT’s inferred gene and protein expressions, we calculated Pearson’s correlation coefficient between the inferred and measured expression values of each cell (cell-wise). Also, we computed the gene-wise correlation between inferred and measured expression values across cells for each gene [10]. For peak inference in open chromatin regions, we used AUROC to evaluate the quality of CMOT’s binarized inferred peaks [10]. We computed peak-wise AUROC between individual inferred peak profiles versus measured profiles. This evaluation also applied to the state-of-art methods that we compared.

#### Classifying known cell type using inferred expression

For the human brain data with known brain cell type information, we evaluated the CMOT inferred expression of cell-type marker genes for classifying the cell type and calculated the AUPRC of the classification [10]. To this end, given a cell type, we labeled all cells that belong to the cell type as positive and the rest as negative. Specifically, we evaluated Top 8 marker genes from each cell type, due to disproportionate cell-type distribution within the dataset, using a total of 80 cells. We then defined a baseline = 0.1 for the AUPRC as the ratio of the number of positives versus total cells.

#### Clustering cancer types using inferred gene expression by Silhouette Score

For the pan cancer dataset, we evaluated CMOT to separate the cancer types. In particular, we assessed if CMOT’s inferred gene expression data can cluster the cells and cell clusters correspond to different cancer types [25], using the Silhouette score:

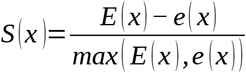
where *E* (*x*) is the average distance between the cell *x* and the other cells of the same cluster and *e* (*x*) is the average distance between *x* and cells in the closest different clusters. We calculated the Silhouette scores by the Python package scikit-learn [23].

#### Gene set enrichment analysis

We used Metascape [26] to perform gene set enrichment analysis for the highly predictive genes by CMOT.

#### Comparison with state-of-arts

We compared CMOT with existing state-of-art methods, Seurat [6], MOFA+ [7], TotalVI [4], and Polarbear [10]. We reported the number of genes with improved correlation/AUROC w.r.t. these methods along with a one-sided Wilcox rank-sum test p-value for each [10]. Unless otherwise stated, we use the term CMOT for the trained model with full correspondence (*p*=100%).

## 3 Results

### 3.1 Single-cell gene expression inference from chromatin accessibility in human and mouse brain

#### Human Brain

We first applied CMOT to single-cell human brain data with jointly profiled chromatin accessibility and gene expression by 10xMultiome (scATAC-seq and scRNA-seq of 8,981 cells) and inferred gene expression of cells from open chromatin regions (OCRs by peaks from scATAC-seq) [1]. We selected top 1000 most variable genes & peaks (Supplementary Methods). We randomly split the cells into 80% training for cross-validation and 20% testing set for evaluation. We split the training set into training and validation to find optimal parameters for the model using 5-fold cross-validation. For the alignment, we set *K*=5, and latent dimension *d*=20. For optimal transport, we set parameters λ=200 and η = 1. For KNN modality inference, we set *k*=600. Also, we used 10 major brain cell types from the dataset. However, to test CMOT’s perfomance when such cell-type information is absent, we induced cell labels by two major cell clusters. We also tested CMOT’s performance for different levels of correspondences: *p*=25%, 50%, 75%, 100%.

CMOT achieves a strong performance for gene expression inference on the testing data, outperforming state-of-the-art methods like Seurat and MOFA+ (**Fig. 2A, 2B**). For instance, CMOT reports a median cell-wise Pearson correlation *r* = 0.67 for *p*=100%, significantly higher than both MOFA+(median *r* = 0.4, Wilcoxon rank-sum test p-value = 0) and Seurat (median *r* = 0.64, Wilcoxon p-value < 1.23e-14). Even for partial correspondences, CMOT has significantly higher performances (median r =0.65 for p=75%, and r=0.65 for p= 50%) than MOFA+ (Wilcoxon p-value < 2.8e-294) and Seurat (Wilcoxon p-value <3.43e-10). Also, with low correspondence such as *p*=25%, CMOT’s performance is still significantly higher than MOFA+ (Wilcoxon p-value < 1.65e-157). For gene-wise correlation (**Fig. 2B**), CMOT *p*=100% and *p*=75%, both outperform MOFA+ for 836 versus 118 genes (Wilcoxon p-value < 5.38e-118) and 827 versus 140 genes (Wilcoxon p-value < 1.78e-165), respectively (**Fig. 2B**). Also, CMOT *p*=100% outperforms Seurat for 494 versus 460 genes (Wilcoxon p-value < 2.42e-2). For CMOT *p*=75%, Seurat slightly performs better for 497 genes versus 471 for CMOT (Wilcoxon p-value < 3.01e-1).

**Figure 2:**
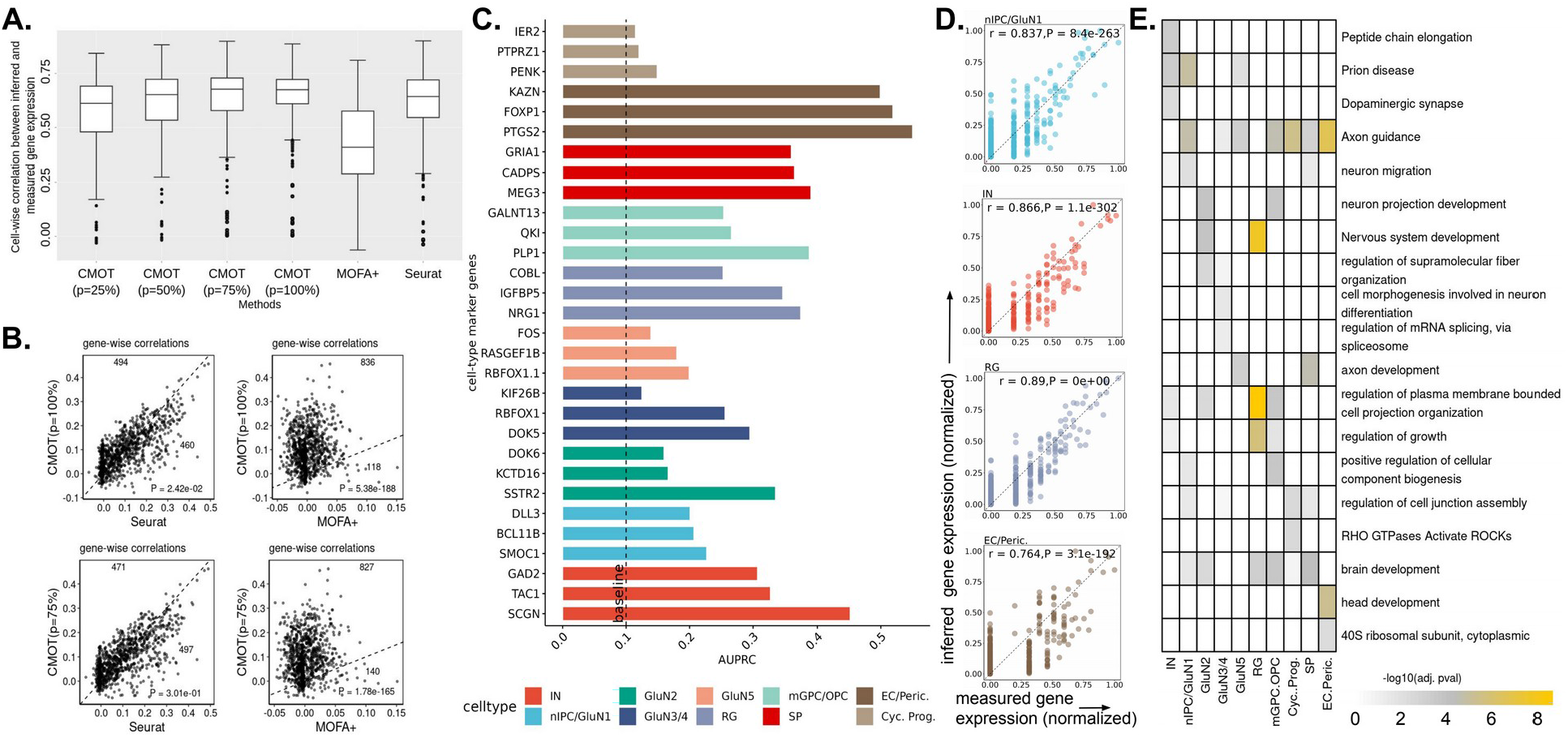
Single-cell gene expression inference from chromatin accessibility in developing human brain. **(A)** Cell-wise Pearson correlation (y-axis) of inferred and measured gene expression by different methods (x-axis): CMOT (*p*=25%,50%,75%,100%), Seurat, MOFA+ (**Table S1**). **(B)** Gene-wise correlation between the inferred and measured gene expression, comparing CMOT (y-axis) with MOFA+ and Seurat (x-axis). Dots: Genes; Numbers: numbers of genes with improved inference by comparing methods. P-values are from onesided Wilcoxon rank-sum tests. **(C)** Gene-wise AUPRC of cell type marker gene for classifying the cell type. Dashline: baseline=0.1 (see Methods). **(D)** The measured (x-axis) versus inferred normalized expression (y-axis) of genes (dots) for one cell across four cell types: nIPC/GluN1(first), E.C./Peric. (second), IN (third), R.G. (last). **(E)** Heatmap showing different enriched terms ranked by -log10(adj. pvalue of enrichment) values for the top 100 highly predictive genes within each cell type (see Methods). *r* is Pearson correlation coefficient. *p* is the correlation p-value.

Next, we evaluated if the CMOT inferred gene expression to classify brain cell types. We used known celltype marker genes provided along with the dataset and selected the top 8 highly predictive cells from each cell type within our inferred gene expression. We then calculated the AUPRC of the respective inferred genes in a one-vs-all manner for each cell type against the rest (Methods). CMOT obtains the higher AUPRCs for these genes against a *baseline* of 0.1 (**Fig. 2C**) for all cell types. The *baseline* is defined as the proportion of positives in the data. This suggests that the CMOT inferred expression is capable to distinguish cell types, providing the biological interpretability of the CMOT inference. Looking at individual cells (**Fig. 2D**), CMOT infers individual cell expression with high Pearson correlation and significance (p<0.05). Furthermore, we also found that the enriched functions and pathways relating to brain development from the top 100 highly predictive genes (Fig. 2E).

#### Mouse Brain

Next, we applied CMOT to a SNARE-seq dataset that jointly profiled gene expression and open chromatin regions of single cells in the adult mouse brain [24]. We also inferred gene expression from peak signals on open chromatin regions and compared CMOT with Polarbear [10] which previously used the same dataset. We split the dataset randomly into training, validation, and testing sets following Polarbear (using its source code). We trained both variation versions of the Polarbear model (Polarbear and Polarbear co-assay) with default parameters. We trained CMOT with the following parameters: *K*=5, *d=* 20, λ=1e03, η = 1e-2, *k*=400, and used the top 10 features of scATAC-seq training data to find the *k* nearest neighbors. We induced our cell labels by hierarchical clustering of the cells in the training set and identified two clusters to regularize the optimal transport in CMOT. We evaluated CMOT and Polarbear by both cell-wise and gene-wise correlations across normalized inferred gene expression (**Fig. S1**). We found that CMOT has significantly higher gene-wise correlations than both Polarbear (515 genes versus 485 genes, Wilcoxon p-value < 2.81e-02) and Polarbear co-assay (648 genes versus 352 genes, Wilcoxon p-value <1.2e-27).

### 3.2 Inferring protein expression from gene expression in peripheral blood mononuclear cells

We applied CMOT to infer protein expression from gene expression of peripheral blood mononuclear cells (PBMCs) using emerging CITE-seq data [4]. We trained CMOT on 6885 cells from PBMC10k, with parameters: *K*=5, *d*= 15, λ=1e02, η = 1, *k*=100, and used the top 200 highly variable genes in the training data to find the *k* nearest neighbors. We induced cell labels by identifying two clusters using gene expression for the label regularization in optimal transport. We evaluated CMOT, MOFA+, Seurat, and TotalVI’s using 3994 cells from a different dataset, PBMC5k. As shown in Fig. 3A, CMOT achieves a median cell-wise Pearson correlation=0.86 for *p*=100%, significantly outperforming MOFA+ (Wilcoxon p-value < 6.9e-57) and TotalVI (default parameters) (Wilcoxon p-value 0) as well as comparable with Seurat. For instance, we show two cells and their Pearson correlation of inferred versus measured protein expressions (*r*=0.99, *p*=1.4e-11 and *r*=0.98, *p*=6.1e-11) in Fig. 3B. Moreover, even for partial correspondences, *p* = 25%, 50%, 75%, CMOT performs consistently with significantly higher cell-wise correlation than MOFA+ (Wilcoxon p-values 0,0,0) and TotalVI (Wilcoxon p-values < 8.36e-58, 1.73e-45, 5.25e-12). Also, for inferring individual protein expression, CMOT has high correlations for all proteins, consistent with state-of-art methods (Fig. 3C), with some examples shown in Fig. 3D. Rest of the proteins along their inference statistics are reported in Fig. S2 and Table S6.

**Figure 3:**
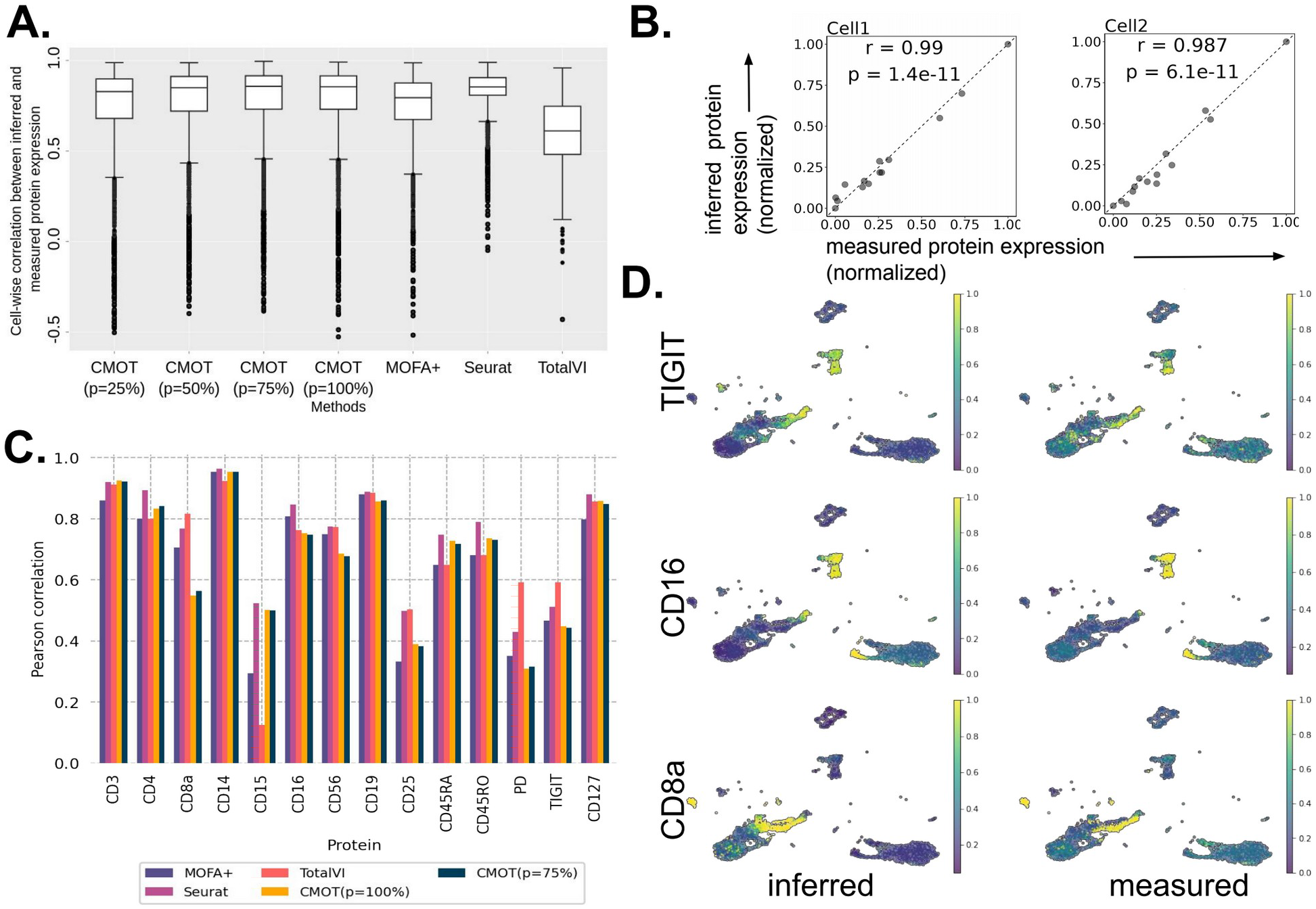
Inferring protein expression from gene expression in single-cell peripheral blood mononuclear cells. **(A)** Cell-wise Pearson correlation (y-axis) of inferred and measured protein expression by different methods (x-axis): CMOT(*p*=25%,50%,75%,100%), Seurat, MOFA+, TotalVI (**Table S2**) **(B)** The measured (x-axis) versus inferred normalized expression (y-axis) of 14 proteins (dots) for two select cells. **(C)** Pearson correlations of inferred and measured expression (y-axis) of individual proteins (x-axis) by CMOT(*p*=100%,75%), Seurat, MOFA+, TotalVI **(D)** UMAPs of inferred and measured expressions for three proteins: TIGIT (r=0.45; p=6.18e-197) (top), CD16 (r=0.75; p=0) (middle), CD8a (r= 0.55; p=3.11e-312) (bottom) (**Table S6**). The intensity represents the protein expression level. r is Pearson correlation coefficient. p is the correlation p-value.

### 3.3 Inference of gene expression using chromatin accessibility for drug-treated lung cancer cells

Next, we applied CMOT to 100nM dexamethasone (DEX)-treated A549 single-cells from lung adenocarcinoma. The cells were profiled after 0, 1, and 3 h of treatment for gene expression and open chromatin regions (OCRs) using sci-CAR experiments [2]. We focus on the CMOT’s performance for gene expression inference from peak signals of OCRs. We stratified-split the dataset into 80% training and 20% test cells using the treatment hours. We used the treatment hours as the classes for label regularization in optimal transport for training cells. We trained CMOT with the parameters: *K*=5, *d=* 10, λ=1e02, η = 1e-3, *k*=500, and used the top 20 highly variable OCRs in scATAC-seq to find the *k* nearest neighbors. Again, we found that CMOT shows a consistent performance across different cell-to-cell correspondence information (*p*) with high correlation. CMOT (*p*=100%) infers gene expression with a median Pearson correlation of 0.52, similar to MOFA+ and significantly outperforming Seurat (median correlation = 0.41, Wilcoxon p-value < 2.19e-57) (**Fig. 4A**). Moreover, CMOT shows a high gene-wise Pearson correlation outperforming Seurat for 636 versus 547 genes (**Fig. 4B**, **Table S7**). Although MOFA+ reports a higher gene-wise Pearson correlation for some genes than CMOT (**Fig. 4B, Table S7**), we still see that CMOT’s inferred expression shows the transitory trend of key druggable marker genes across drug-treatment hours. **Fig. 4C** shows three key genes, identified as makers of early (ZSWIM6) [30] and late (PER1, BIRC3) [27]–[29] events of treatment. Also, we performed enrichment of the 435 high correlation genes identified by CMOT (versus MOFA+ in **Fig. 4B**). As shown in **Fig. 4D**, we saw a higher enrichment of terms associated with DEX-treated A549 cells like TGF-beta signaling, along with effects on DEX treatment in general, like Mental disorders as compared to enrichment given by MOFA+ (**Fig. S3**). Lastly, we also found that the cell-wise correlations between inferred and measured gene expression are also significantly highly correlated in each treatment hour (**Fig. 4E**).

**Figure 4:**
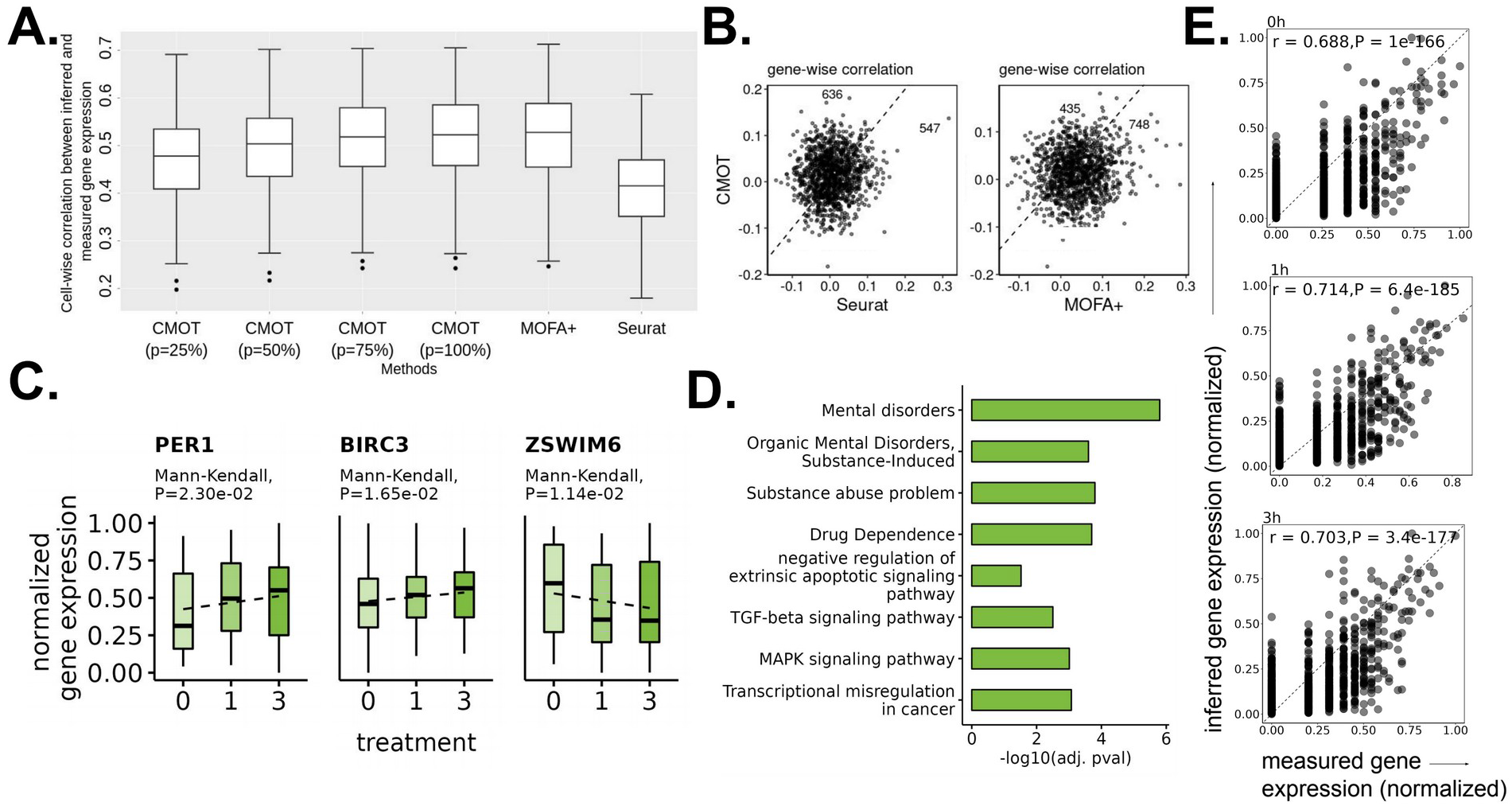
Inference of gene expression for drug-treated lung cancer cells using chromatin accessibility. **(A)** Cell-wise Pearson correlation (y-axis) of inferred and measured gene expression by different methods (x-axis): CMOT(*p*=25%,50%,75%,100%),Seurat, MOFA+ (**Table S3**) **(B)** Gene-wise correlation between the inferred (y-axis) and measured (x-axis) expression, comparing CMOT with MOFA+ and Seurat. Dots: Genes; Numbers: Gene numbers above and below the dotted line. P-values are calculated by a one-sided Wilcoxon rank-sum test (**Table S7**) **(C)** CMOT inferred normalized gene expression trend (y-axis) across treatment hours (x-axis). Key genes: PER1 and BIRC3 [27]–[29] are markers for glucocorticoid receptor (GR) activation seen later in treatment (3h). ZSWIM6 [30] is a key gene of early events of DEX treatment (0h,1h) **(D)** Enriched terms associated with CMOT inferred gene expression using 435 genes with a higher gene-wise Pearson correlation compared to MOFA+’s 748 genes inference in (**B, D, Fig. S3**) **(E)** The measured (x-axis) versus inferred normalized expression (y-axis) of genes (dots) for three select cells. r is Pearson correlation coefficient. p is the correlation p-value.

### 3.4 Cross-modality inference between gene expression and chromatin accessibility to distinguish cancer types

Finally, we tested CMOT to see how well it can infer between two modalities, especially for relevantly small datasets. We used a pan-cancer scCAT-seq dataset which jointly profiled single-cell gene expression and chromatin accessibility on OCRs for three cancer cell lines: HCT116, HeLa-S3, and K562 [25]. We stratified split data into 60% training and 40% testing sets using cancer-type information. We induced our own cell labels for training cells for label regularization in optimal transport. For gene expression inference from OCR peaks, we identified two clusters in chromatin peaks and vice versa. We trained CMOT with the following parameters for gene expression inference from chromatin peaks: *K*=5, *d=* 10, λ=1e03, η = 1, *k*=40, and used the top 150 highly variable OCRs to find the *k* nearest neighbors. For inferring gene expression from binarized OCR peaks, we evaluated the inferred expression using the same metrics (cell-wise and gene-wise Pearson correlation) as above. CMOT significantly outperforms both MOFA+ and Seurat, with a cell-wise median correlation of 0.69 compared to 0.65 (Wilcoxon p-value < 6.81e-17) and 0.51 (Wilcoxon p-value < 1.32e-05), respectively (**Fig. 5A, Table S4**). Moreover, CMOT (*p*=100%) yields an improved gene-wise correlation for 6307 genes versus 3693 against Seurat (Wilcoxon p-value < 7.19e-18), and, 8734 versus 1266 against MOFA+ (Wilcoxon p-value = 0) (**Fig. 5D**). Moreover, CMOT’s inference is particularly useful to identify the cancer type speciifc cell clusters. For instance, we calculated the Silhouette score (see Methods) to see if the cells from the same cancer lines exhibit similar gene expression patterns. CMOT reports a high median silhouette score of 0.82 compared to the measured gene expression (0.35) and inferred expressions from Seurat (0.75) and MOFA+ (0.05) (**Fig. 5B, Table S5**). As shown in **Fig. 5C**, the cancer cells from three cancer cell lines can be separated using CMOT inferred gene expression, suggesting the capability of CMOT inference to reveal cancer-type-specific expression. Then, we evaluated the CMOT’s OCR peaks inference from gene expression. We trained CMOT with the parameters: *K*=5, *d=* 10, λ=1e03, η = 1, *k*=10, and used the top 50 highly variable genes to find the *k* nearest neighbors. We also stratified split the data into 60% training and 40% testing sets using cancer-type information. We normalized CMOT’s inferred peaks and then binarized them by a cutoff 0.5, and then calculated the peak-wise area under the receiver operating curve (AUORC) of the inferred binarized peaks relative to the binarized measured profile. We also found that CMOT significantly outperforms both MOFA+ and Seurat with Wilcoxon p-values < 2.71e-86, 2.70e-72 respectively for OCR peak inference from gene expression (**Fig. 5E, Fig. S4, Table S9**).

**Figure 5:**
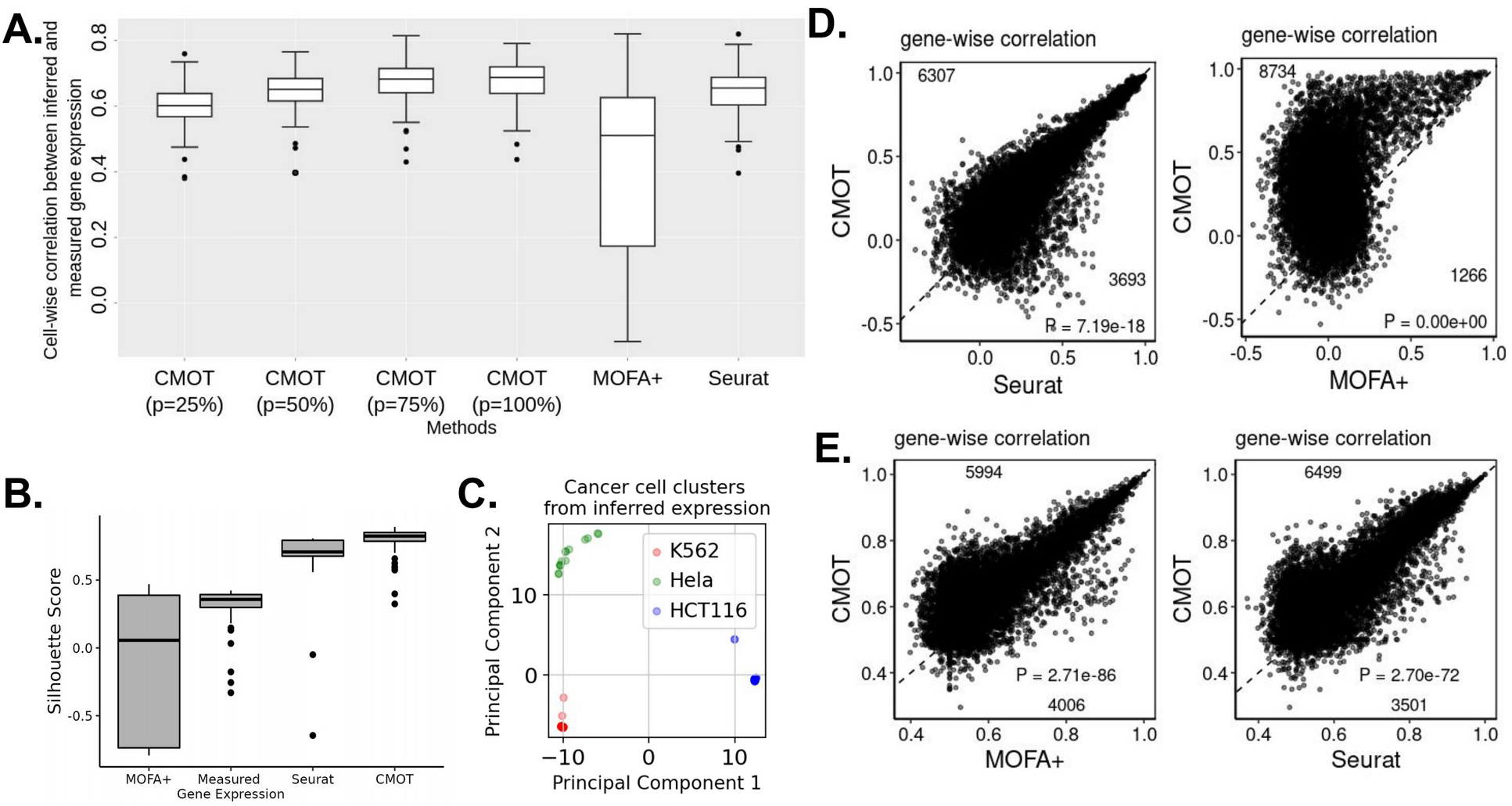
Cross-modality inference between gene expression and chromatin accessibility can distinguish cancer types. **(A)** Cell-wise Pearson correlation (y-axis) of inferred and measured gene expression by different methods (x-axis): CMOT(*p*=25%,50%,75%,100%),Seurat, MOFA+ (**Table S4**) **(B)** Silhouette score (x-axis) across measured and inferred gene expressions (x-axis) (**Table S5**) **(C)** PCA of inferred gene expression **(D)** Gene-wise correlation between the inferred and measured expression, comparing CMOT (y-axis) with MOFA+ and Seurat (x-axis). Dots: Genes; Numbers: Gene numbers above and below the dotted line **(E)** Peak-wise AUROC, comparing CMOT (y-axis) with MOFA+ and Seurat (x-axis). Dots: Peaks; Numbers: Peak numbers above and below the dotted line. P-values are calculated by a one-sided Wilcoxon rank-sum test.

## 4 Discussion

CMOT is a computational approach that integrates manifold alignment, regularized optimal transport, and k-nearest neighbors (KNN) for cross-modality inference. By applying emerging single-cell multimodal data, we demonstrated that CMOT was able to predict multimodal features of single cells such as gene expression, chromatin accessibility, and protein expression. Note that CMOT does not require paired samples for aligning multimodalities as shown by its outperformances over state-of-the-arts in some applications. This attribute is particularly useful since joint multimodal profiling is typically challenging and sometimes costly and single modality data is thus still widely favored. In the paper, CMOT primarily used nonlinear manifold alignment (NMA) to align multimodalities for achieving the best inference. However, CMOT is flexible and the user can substitute NMA with their preferred alignment method, e.g., SCOT [13], MMD-MA [31], and WNN [6].

Furthermore, the optimal transport step in CMOT leverages the information within shared modalities between source and target cells to compute a mapping matrix through Wasserstein distances between them. These distances quantify and minimize the geometric discrepancy of the distributions and map the two distributions for improving cross-modal inference in CMOT.

Nonetheless, we also examined CMOT’s potential limitations. First, nonlinear manifold alignment comes with a high computational cost as the data size increases. It needs to compute Laplacians and similarity matrices of multimodal inputs which scale quadratically with the datasets and slows CMOT’s computations over large datasets (**Table S10**). However, this can potentially be sped up by using other alignment methods like SCOT [13], [14] or Unioncom [32]. Second, Optimal transport assumes a mass-balancing approach between the source and target distributions, where every s (e.g., a cell) in the source has to map to a point in the target. This is a relatively strong assumption requiring a balanced data distribution between the source and target to fit a conservative transport plan. Given many imbalanced datasets in the real world, this limitation can be improved by recent optimal transport techniques such as SCOTv2 [14] through emerging unbalanced optimal transport approaches [33], [34].

Also, CMOT can adapt other optimal transport variants to even transport different modalities between the source and target cells. For instance, Gromov Wasserstein distance can map distributions from different modalities [35], [36]. Moreover, CMOT has the potential to work with additional single-cell modalities like morphology of single neuronal cells from Patch-seq. In addition to inferring modalities, CMOT has the potential to be extended to infer the sample labels such as phenotypes across modalities, e.g., via label transferring [6]. For example, it can predict cell types or disease states of single cells for the modalities without such information.

## 5 Acknowledgments data and code availability, fundings and supplementary files

This work was supported by National Institutes of Health grants, R01MH128695-01A1, R21NS128761, R21NS127432, R01AG067025, R21CA237955, R01HD106197, U01MH116492, R03NS123969 and National Science Foundation Career Award 2144475. All data and code are available at https://github.com/daifengwanglab/CMOT. The supplementary tables, figure and notes are provided in Supplementary materials.

## Notes

### Competing Interest Statement

The authors have declared no competing interest.

