## Supplementary Materials for "CMOT: Cross Modality Optimal Transport for multimodal inference"

### 1 Supplementary Tables

| **Methods** | **Seurat** | **MOFA+** |
|---|---|---|
| **CMOT (*p*=100%)** | 1.23×10−14 | 0 |
| **CMOT (*p*=75%)** | 3.4×10−10 | 2.8x10-294 |
| **CMOT (*p*=50%)** | 0.3 | 3.3x10-240 |
| **CMOT (*p*=25%)** | 1 | 1.65x10-157 |

**Table S1: Gene expression inference from chromatin accesibilty in human brain data [1].** Wilcox-rank sum test p-values of cell-wise Pearson correlation between inferred and measured gene expression comparing methods (x-axis) CMOT(*p*=25%,50%,75%,100%), Seurat, and, MOFA+ plotted in Fig. 2A for human brain [1]. All values have been calculated between CMOT variations (*p*=100%,75%,50%,25%) , and competing methods by setting alternative=” greater”.

| **Methods** | **Seurat** | **MOFA+** | **TotalVI** |
|---|---|---|---|
| **CMOT (*p*=100%)** | 1 | 6.9 × 10−57 | 0 |
| **CMOT (*p*=75%)** | 1 | 0 | 8.36×10−58 |
| **CMOT (*p*=50%)** | 1 | 0 | 1.73×10−45 |
| **CMOT (*p*=25%)** | 1 | 0 | 5.25×10−12 |

**Table S2: Protein expression inference from gene expression in PBMC [4].** Wilcox-rank sum test p-values of cell-wise Pearson correlation between inferred and measured gene expression comparing methods (x-axis) CMOT(*p*=25%,50%,75%,100%), Seurat, and, MOFA+ plotted in Fig. 3A for Peripheral Blood Mononuclear Cells (PBMCs) [4]. All values have been calculated between CMOT variations (*p*=100%,75%,50%,25%) , and competing methods by setting alternative=” greater”.

| Methods | Seurat | MOFA+ |
|---|---|---|
| **CMOT (*p*=100%)** | $2.19\times10^{-57}$ | 0.65 |
| **CMOT (*p*=75%)** | $2.34\times10^{-55}$ | 0.81 |
| **CMOT (*p*=50%)** | $3.95\times10^{-43}$ | 0.99 |
| **CMOT (*p*=25%)** | $2\times10^{-24}$ | 1 |

**Table S3: Gene expression inference from chromatin accessibility in DEX-treat A549 lung cancer data [2].** Wilcox-rank sum test p-values of cell-wise Pearson correlation between inferred and measured gene expression comparing methods (x-axis) CMOT (*p*=25%,50%,75%,100%), Seurat, and, MOFA+ plotted in Fig. 4A for DEX-treated A549 cells [2]. All values have been calculated between CMOT variations (*p*=100%,75%,50%,25%) , and competing methods by setting alternative="greater".

| Methods | Seurat | MOFA+ |
|---|---|---|
| **CMOT (*p*=100%)** | $1.32\times10^{-05}$ | $6.81\times10^{-17}$ |
| **CMOT (*p*=75%)** | $7.88\times10^{-05}$ | $3.68\times10^{-16}$ |
| **CMOT (*p*=50%)** | 0.4 | $1.07\times10^{-11}$ |
| **CMOT (*p*=25%)** | 0.99 | $7.42\times10^{-06}$ |

**Table S4: Gene expression inference from chromatin accessiblity in pan-cancer data [25].** Wilcox-rank sum test p-values of cell-wise Pearson correlation between inferred and measured gene expression comparing methods (x-axis) CMOT (*p*=25%,50%,75%,100%), Seurat, and, MOFA+ plotted in Fig. 5A for pan-cancer cells [25]. All values have been calculated between CMOT variations (*p*=100%,75%,50%,25%) , and competing methods by setting alternative="greater".

| Methods | Seurat | MOFA+ | Measured Expression |
|---|---|---|---|
| **CMOT** | $2.28\times10^{-15}$ | $2.54\times10^{-28}$ | $5.96\times10^{-28}$ |

**Table S5: Silhouette score across measured and inferred gene expression in pan-cancer data [25].** Wilcox-rank sum test p-values for Silhouette scores reported in Fig. 5B for Pan-cancer cells [25]. All p-values have been calculated between the silhouette scores reported for inferred gene expressions by CMOT (*p*=100%), Seurat, MOFA+, and, measured expression, by setting alternative="greater".

| Protein | Pearson Correlation | P-value |
|---|---|---|
| **CD3** | 0.92 | 0 |
| **CD4** | 0.83 | 0 |
| **CD8a** | 0.54 | 3.1×10−312 |
| **CD14** | 0.95 | 0 |
| **CD15** | 0.5 | 1.58×10−252 |
| **CD16** | 0.75 | 0 |
| **CD56** | 0.69 | 0 |
| **CD19** | 0.86 | 0 |
| **CD25** | 0.4 | 3.06×10−144 |
| **CD45RA** | 0.72 | 0 |
| **CD45RO** | 0.73 | 0 |
| **PD-1** | 0.31 | 2.36×10−89 |
| **TIGIT** | 0.45 | 6.17×10−197 |
| **CD127** | 0.85 | 0 |

**Table S6: Pearson correlation between inferred and measured protein expression in PBMC [4]:** Pearson correlation and p-values between CMOT's inferred and measured protein expression for PBMCs 5K [4] (Fig. 3B,C)

| Methods (m) | #genes (CMOT>m) | #genes(CMOT<m) | P-value (CMOT>m) |
|---|---|---|---|
| **MOFA+** | 636 | 547 | 2.19×10−06 |
| **Seurat** | 435 | 748 | 1 |

**Table S7: Gene-wise correlation between inferred and measured gene expression in DEX-treat A549 lung cancer data [2].** Number of genes with higher gene-wise Pearson correlation between inferred and measured gene expression profiles for DEX-treated A549 lung adenocarcinoma [2] cells. Column1 reports the number of genes inferred by CMOT that have a higher correlation than the respective method m. Column2 reports vice versa of column1. Column3 reports the p-value statistic for the Wilcox-rank sum test between gene-wise correlations of CMOT and competing methods (alternative="greater").

| Methods | Polarbear | Polarbear-coassay |
|---|---|---|
| **CMOT (*p*=100%)** | 0.79 | 0.29 |
| **CMOT (*p*=75%)** | 0.79 | 0.29 |
| **CMOT (*p*=50%)** | 0.99 | 0.99 |
| **CMOT (*p*=25%)** | 1 | 1 |

**Table S8: Gene expression inference from chromatin accessiblity in mouse brain [23].** Wilcox-rank sum test p-values of cell-wise Pearson correlation between inferred and measured gene expression comparing methods (x-axis) CMOT (*p*=25%,50%,75%,100%), Polarbear, and, MOFA+ plotted in Fig. S1 for mouse brain [23]. All values have been calculated between CMOT variations (*p*=100%,75%,50%,25%) , and competing methods by setting alternative="greater".

| Methods | Seurat | MOFA+ |
|---|---|---|
| **CMOT (*p*=100%)** | 0.18 | 0.73 |
| **CMOT (*p*=75%)** | 0.09 | 0.59 |
| **CMOT (*p*=50%)** | 0.32 | 0.8 |
| **CMOT (*p*=25%)** | 0.99 | 0.99 |

**Table S9: chromatin accessibility inference from gene expression in pan-cancer data [25].** Wilcox-rank sum test p-values of cell-wise Pearson correlation between inferred and measured gene expression comparing methods (x-axis) CMOT (*p*=25%,50%,75%,100%), Seurat, and, MOFA+ plotted in Fig. 5A for pan-cancer cells [25]. All values have been calculated between CMOT variations (*p*=100%,75%,50%,25%) , and competing methods by setting alternative="greater".

| Methods | Developing Human Brain[1] | Mouse Brain [23] | PBMCs [4] | DEX-treated A549 [2] | Liu Cancer cell lines [25] (gene expression from chromatin accessibility) | Liu Cancer cell lines [25] (chromatin accessibility from gene expression ) |
|---|---|---|---|---|---|---|
| **CMOT** | 1810 | 673.54 | 2038 | 42.13 | **3.34** | **2.19** |
| **Seurat** | **49.74** | - | **347.05** | **18.35** | 11.46 | 13.61 |
| **MOFA+** | 264.13 | - | 1953.2 | 272.3 | 226.54 | 164.91 |
| **TotalVI** | - | - | 426.36 | - | - | - |
| **Polarbear** | - | 964 | - | - | - | - |
| **Polarbear-coassay** | - | **292** | - | - | - | - |

**Table S10**: **Running times (in seconds) of all methods across datasets.** (See supplementary Runtime evaluations)

**2 Supplementary Figures**

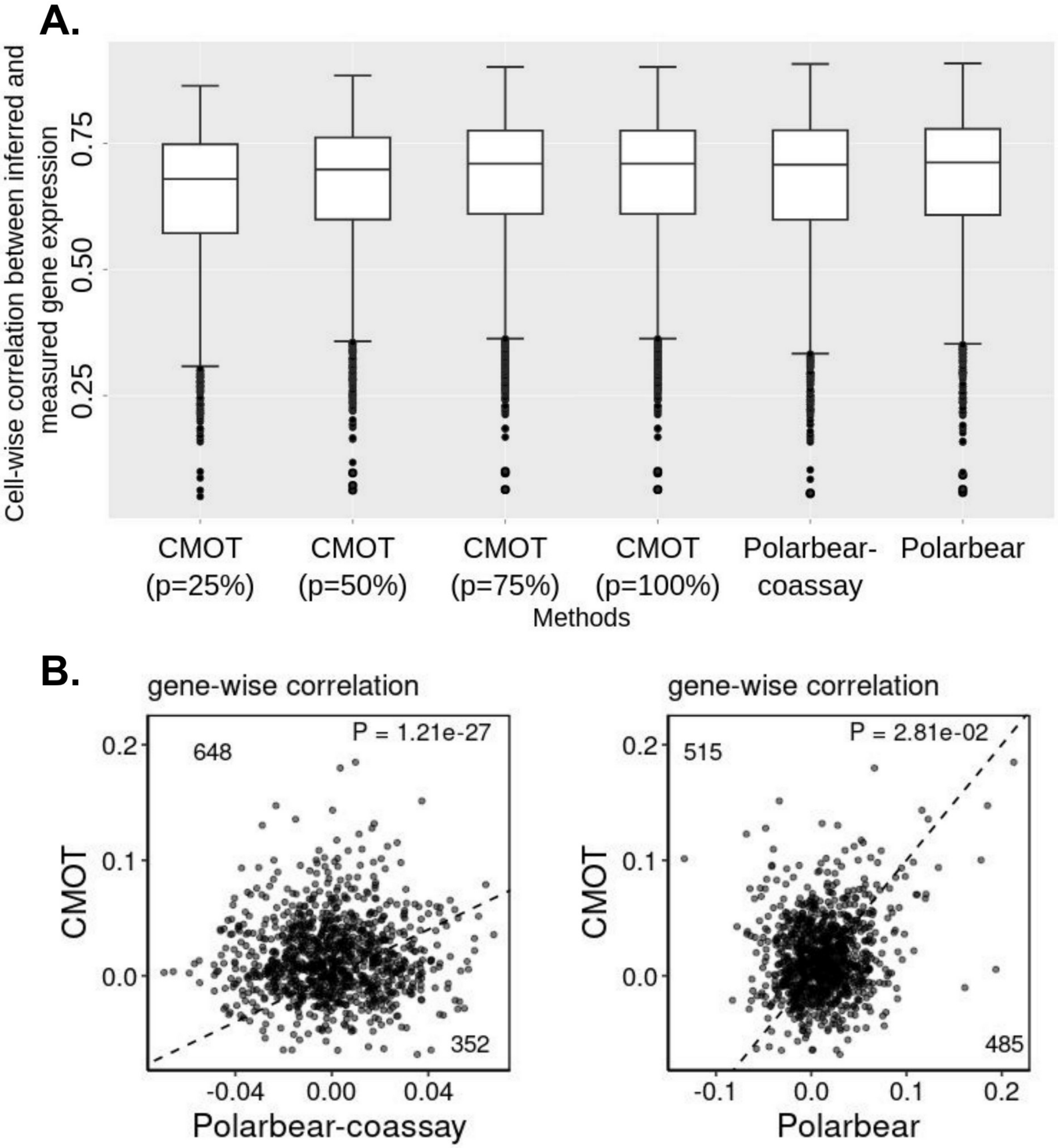


**Figure S1: Gene expression inference from chromatin accessibility in mouse brain (A)** Cell-wise Pearson correlation (y-axis) of inferred and measured gene expression by different methods (x-axis): CMOT(*p*=25%,50%,75%,100%), Polarbear co-assay, and Polarbear (**Table S8**) **(B)** Gene-wise correlation between the inferred and measured gene expression, comparing CMOT (y-axis) with MOFA+ and Seurat (x-axis). Dots: Genes; Numbers: numbers of genes with improved inference by comparing methods. P-values are from one-sided Wilcoxon rank-sum tests.

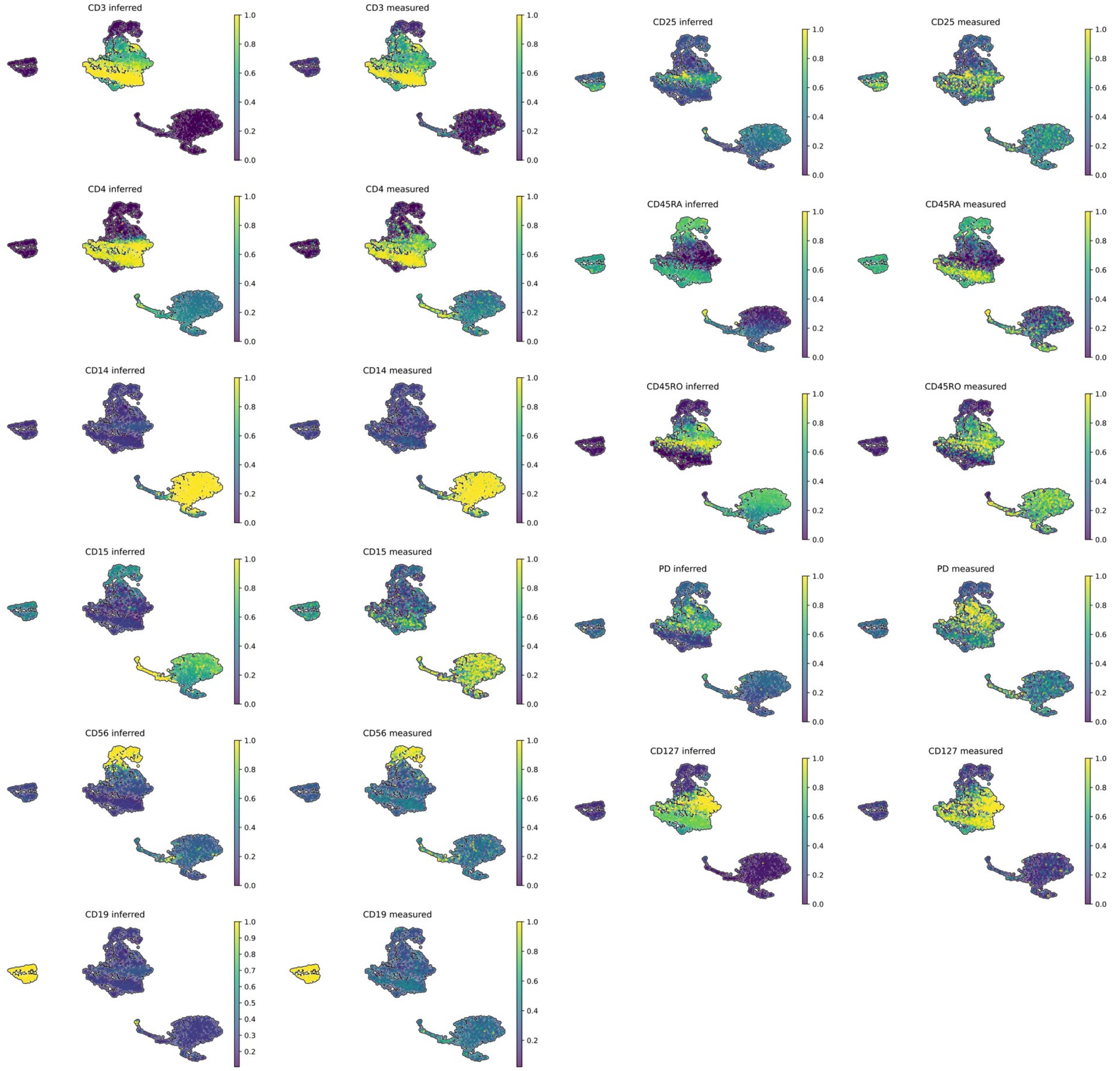


**Figure S2: Inferring protein expression from RNA in single-cell peripheral blood mononuclear cells.** Inferred versus Measured proteins expressions for Peripheral Blood Mononuclear Cells (PBMCs) [18] by CMOT. Included as supplementaryFigure3.pdf with supplementary files.

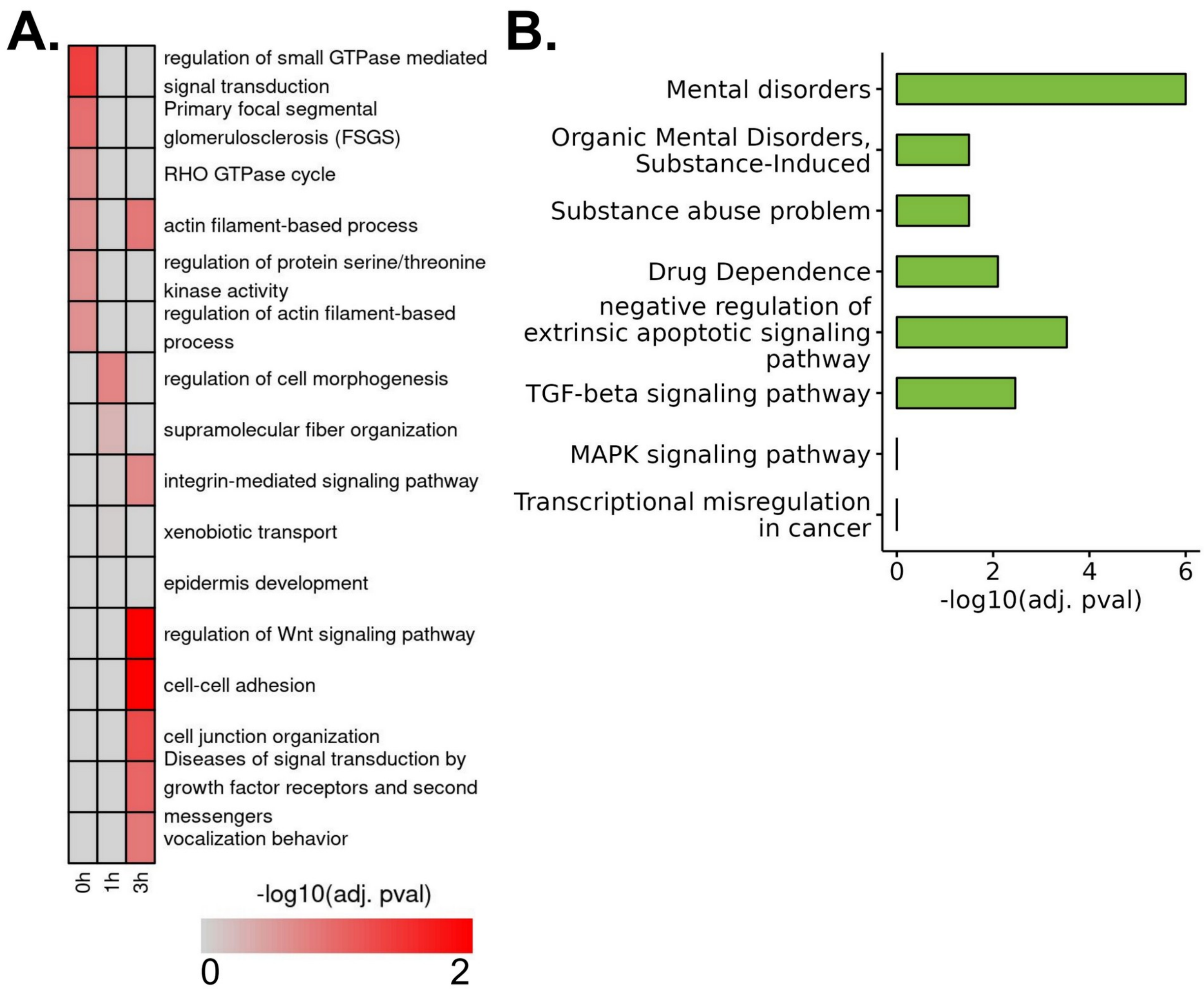


**Figure S3: Inference of gene expression for drug-treated lung cancer cells using chromatin accessibility. (A)** Heatmap showing different enriched terms ranked by -log10(adj. pvalue of enrichment) values for the top 100 highly predictive genes within each treatment hour (see Methods) **(B)** Enriched terms associated with MOFA+ inferred gene expression using 748 genes with a higher gene-wise Pearson correlation compared to CMOT's 435 genes inference in (**Fig. 4B, D**).

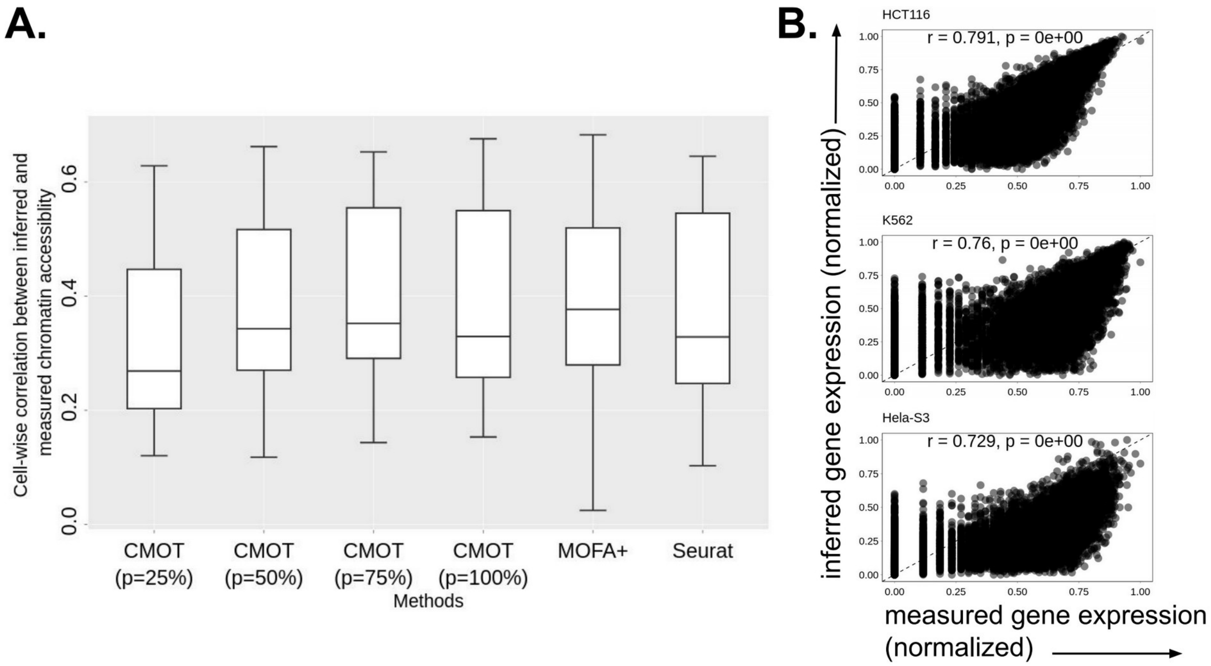


**Figure S4: Cross-modality inference between gene expression and chromatin accessibility in pan-cancer cells (A)** Cell-wise Pearson correlation (y-axis) of inferred and measured chromatin accessibility by different methods (x-axis): CMOT (*p*=25%,50%,75%,100%), Seurat, MOFA+ (**Table S9**) [21]. **(B)** The measured (x-axis) versus inferred normalized expression (y-axis) of genes (dots) for three select cells. r is the Pearson correlation coefficient. p is the correlation p-value.

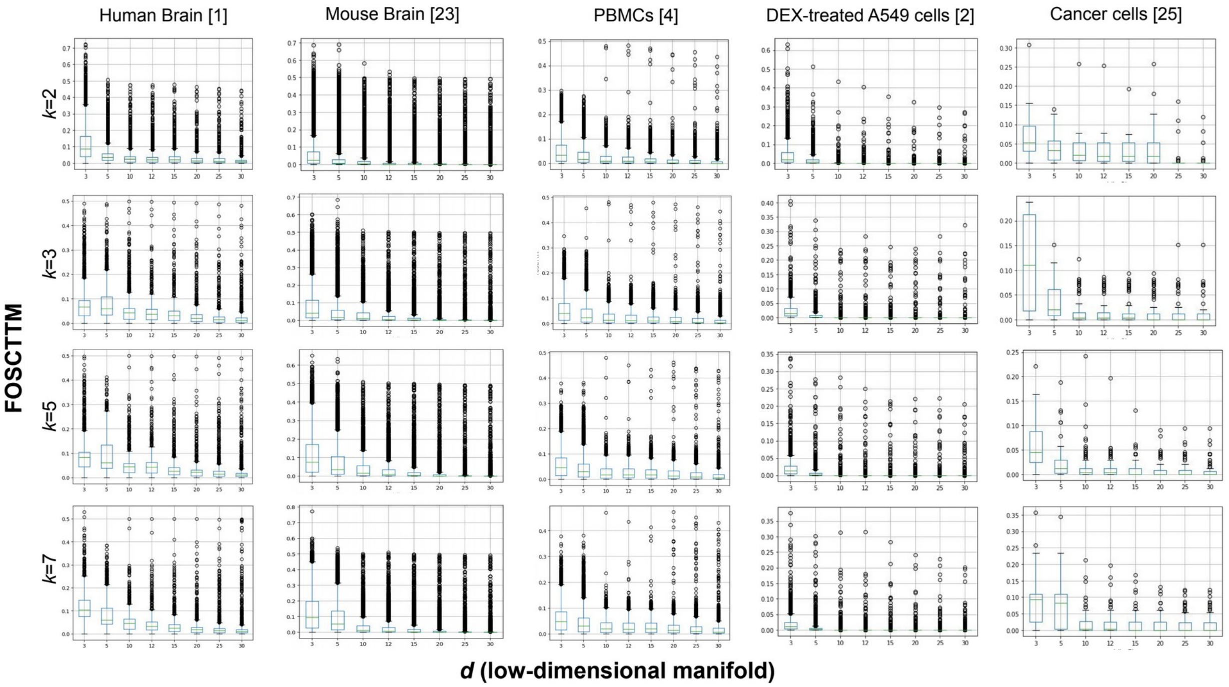


**Figure S5:** Boxplots show the pairwise cell Mean FOSCTTM Score (y-axis) after alignment on the latent dimension *d* (x-axis) for different choices of *k* nearest neighbors (row) in non-linear manifold learning (NMA) for different datasets (column).

### 3 Supplementary Methods

#### 3.1 Datasets preprocessing and feature selection

**Human and mouse brain:** The human brain dataset was generated by 10xGenomics (Gene Expression Omnibus accession GSE162170), containing gene expression and open chromatin regions multiome data from the same cells (8,981 cells) profiled from post-conceptual week 21 (PCW21) [1]. We filtered out peaks and genes that occurred in less than 3 cells. For scATAC, we normalized the peaks using term frequency-inverse document frequency (TF-IDF) transformation using RunTFIDF [39] to identify the top 1000 most variable peaks. For scRNA, we performed normalization and variance stabilization using SCTransform [37] and picked the top 1000 most variable genes. The resulting data includes gene expression and chromatin regions of 8,981 for 1000 genes and regions respectively.

The adult mouse brain dataset [24] was generated by SNARE-seq, containing jointly profiled gene expression and open chromatin regions for ~10k cells. We filtered out peaks and genes that occurred in less than 3 cells. For the binary scATAC data, we picked the top 1000 peaks that were expressed in the largest number of cells. For scRNA, we performed normalization and variance stabilization using SCTransform [37] and picked the top 1000 most variable genes. The resulting data includes gene expression and chromatin regions of 10,839 cells for 1000 genes and regions respectively.

**Peripheral Blood Mononuclear Cells:** The Peripheral Blood Mononuclear cells (PBMC) dataset [4] was generated by CITE-seq, containing genes and proteins from the same cells. This data contains cells from two experiments performed on PBMCs: 6855 cells from PBMC10k and 3994 cells from PBMC5k. We preprocessed multiome data from two experiments independently. For scRNA-seq, performed normalization and variance stabilization using SCTransform [37], and picked 2960 highly variable genes. We identified the variable genes in PBMC10k first and used them as a reference to subset genes in PBMC5k scRNA data. For protein expression, we performed centered log-ratio (CLR) normalization using Seurat's functions in both PBMC10k and PBMC5k. The resulting PBMC10k data dimension was 6855 $X$ 2960 for gene expression and 6855 $X$ 14 for protein expression. The resulting PBMC10k data includes gene and protein expression data of 3994 cells for 2960 genes and 14 proteins.

**Dexamethasone-treated A549 cells:** The dexamethasone(DEX)-treated A549 dataset [2] was generated using sci-CAR experiment for single cells from the A549 lung adenocarcinoma cell line. The data contains jointly profiled 2,641 cells after 0, 1, and 3 h of 100 nM DEX treatment for gene expression and open chromatin regions. We used a preprocessed dataset previously used by Jin et al. [38], and filtered out lowly expressed cells by gene expression. We reduced the dataset to 2391 cells. For scRNA, we used all 1183 genes. For scATAC, we picked the top highly variable 1183 peaks. The resulting data includes gene expression and chromatin regions of 2391 cells for 1183 genes and regions respectively.

**Pan-cancer cell lines:** This dataset contains three cancer cell lines [25]: HCT116, HeLa-S3, and, K562, generated by joint profiling of 206 single cells using scCAT-seq containing gene expression and open chromatin regions. We used a preprocessed dataset previously used by Huizing et al.[16], however, we reduced the number of genes and peaks to 10,000 by selecting the most variable features. We binarized the scATAC profile where we set all values greater than 0 to 1 and 0 otherwise. The resulting data includes gene expression and chromatin regions of 206 cells for 10,000 genes and regions respectively.

#### 3.2 Runtime evaluations

We compare CMOT's running time with state-of-art methods MOFA+, Seurat, TotalVI, and Polarbear for the best-performing parameters used for cross-modality inference (**Table S8**). For smaller datasets i.e. A549 lung cancer cells (sci-CAR) [20] and pan-cancer cells (scCAT-seq) [21], we benchmarked all methods on Intel Core i7-1065G7 CPU @ 1.30GHz with 15.4 GiB RAM. For larger datasets i.e. human and mouse brains (scRNA-seq & scATAC-seq) [19,1], and, peripheral blood mononuclear cells (CITE-seq) [18], we benchmarked all methods on Intel Xeon Gold 6242R CPU @3.10GHz x 40 with 251.4GiB RAM and NVIDIA RTX A6000 GPU.

#### 3.3 Hierarchical Clustering

We induced cell labels for datasets with no prior knowledge (e.g., cell types). We use these labels for label regularization in OT optimization (see Methods and Materials Step B) to improve the mappings between cells in source ($X$) and target ($Y$) modalities. To induce cell labels, we performed hierarchical clustering of training and validation sets combined using the scikit-learn clustering functions [23].

#### 3.4 Training and cross-validation

We split the human brain [1], PBMCs [4] and DEX-A549 lung cancer datasets into 80% train and 20% test. We split the pan-cancer dataset [25] into 60% train and 40% test. For the adult mouse dataset [24], we used the same splits as Polarbear i.e. random splitting into 60% train, 20% validation and 20% test.

We trained all methods: Seurat [6], MOFA+ [7], TotalVI [4] and Polarbear [10] using default parameters. For modality inference in Seurat [6], we integrated the training modalities first, then we inferred the missing modality using FindTransferAnchor and TransferData functions [6] between the integrated training modalities and sorurce test modality. For MOFA+ [7], we input the missing modality as NA values and trained the model on the multimodalities. We trained TotalVI [4] autoencoder with default parameters, with latent distribution set to "normal", on the training set. Finally, we trained Polarbear and Polarbear co-assay models [10] using default parameters on the training set.

To identify the highest performing parameters for Steps B and C of CMOT, we performed 5-fold cross validation on the training set. We reported the best performing parameters for each dataset in Results.
